# Integrated histological and proteomic mapping of pancreatic adaptations during porcine pregnancy

**DOI:** 10.64898/2026.05.19.726186

**Authors:** Christos Karampelias, Susanne Badeke, Christine von Toerne, Mireia Molina van den Bosch, Daniel Veselinovic, Kaiyuan Yang, Eckhard Wolf, Elisabeth Kemter, Heiko Lickert

**Affiliations:** Institute of Diabetes and Regeneration Research, Helmholtz Munich, Neuherberg, Germany; German Center for Diabetes Research (DZD), Neuherberg, Germany; Metabolomics and Proteomics Core, Helmholtz Center Munich, German Research Center for Environmental Health, D-80939 Munich, Germany; School of Medicine and Health, Technische Universität München, Munich, Germany; Gene Center and Center for Innovative Medical Models (CiMM), LMU Munich, Munich, Germany; Interfaculty Center for Endocrine and Cardiovascular Disease Network Modelling and Clinical Transfer (ICONLMU), LMU Munich, Munich, Germany

**Keywords:** Pancreatic pregnancy adaptations, beta-cell expansion, pig model, FFPE proteomics

## Abstract

Pregnancy is a period of extensive metabolic rewiring. Insulin secreting β-cells respond to the metabolic challenges of pregnancy by increasing their mass and size and by altering secretory patterns to maintain glucose homeostasis. If glucose metabolism is not tightly controlled, gestational diabetes may develop. Most studies on β-cell adaptation during pregnancy are derived from rodent models, making translation to the vastly different human gestational setting challenging. In this work, we performed an extensive characterization of pancreatic adaptations throughout porcine pregnancy. Pigs have a long gestational period (114 days) and share a similar size and metabolism to humans, making them an ideal model to bridge the knowledge gap between rodents and humans. By analyzing pancreatic samples from early and late gestational ages, we captured the full trajectory of endocrine remodeling. We observed pregnancy-driven remodeling of endocrine cell types, marked by preferential expansion of pancreatic polypeptide-secreting cells. Proteomic characterization of the pancreas from early and late gestation showed a downregulation of SLC20A2 and ZCCHC7, identifying new protein targets involved in physiological endocrine cell adaptation. Overall, our comprehensive characterization of pancreatic adaptations in the pig model helps bridge the translational gap between rodents and humans and highlights previously unrecognized proteins with therapeutic potential for gestational diabetes.

## Introduction

Human pregnancy is characterized by increasing maternal metabolic demands throughout gestation. One of the main metabolic demands is the need to supply the developing fetus with glucose. To cope with this need, particularly in late gestation, mothers develop peripheral insulin resistance [1,2], which increases glucose supply to the fetus. Failure of adaptive mechanisms can lead to gestational diabetes, which is characterized by persistent maternal hyperglycemia [3]. Gestational diabetes constitutes a higher risk for developing type 2 diabetes later in life [4] and the persistent hyperglycemia can have severe effects on the development of the fetus [5]. On the contrary, pregnant women living with type 1 diabetes have a temporary increase in insulin secretion [6] underlying the profound effect of pregnancy on glucose metabolism. Therefore, it is critical to understand metabolic pregnancy adaptations on the molecular level and design new therapeutics for gestational diabetes management.

The endocrine and hormone-producing part of the pancreas is a tissue undergoing major pregnancy-driven adaptation to metabolic needs. Specifically, research in rodent models has shown drastic adaptive changes of the pancreatic insulin producing β-cells in terms of secretory capacity, mass (hyperplasia) and size (hypertrophy) [7]. It has been demonstrated that new β-cells form predominantly through proliferation of the available β-cell mass [8–12], with few studies pointing to transdifferentiation of α-cells to β-cells [13,14] or transdifferentiation/differentiation of ductal residing cells to β-cells [15–17] as alternative cellular mechanisms of β-cell mass adaptation. Lactogen and serotonin signaling pathways are among the most validated in relation to β-cell proliferation and insulin secretion across gestation in mammals [8,10–12,18,19]. Understanding the cellular and molecular mechanisms of the β-cell mass expansion under insulin resistance, as the one developing in pregnancy, can provide new pathways for stimulating β-cell regeneration as a curative approach for diabetes [20].

Given the scarcity of pancreas tissue from cadaveric donors during gestation, few studies have tried to translate the rodent findings to human biology. As it was demonstrated in histological analysis, human β-cells do not undergo such dramatic expansion during gestation with lower levels of proliferation being observed [21–23]. Moreover, lactogen signaling appears to have evolutionary distinct roles in islet biology [18]. Lastly, extensive human studies on the non β-cell islet populations or non-endocrine populations by focusing on distinct developmental timepoints are lacking [7]. A recent FFPE proteomics characterization of endocrine and exocrine human pancreas during pregnancy did not show a pronounced phenotype[23], suggesting that pregnancy-driven pancreas adaptations might be less profound than the ones suggested in rodents on the molecular level[24,25]. Therefore, there is a need for human-relevant models to translate gestational pancreas biology to humans.

In this work, we characterized the cellular and molecular adaptations of the pancreas during pig gestation. Pigs have an extended gestation time (114 days) compared to rodents and their metabolism, islet development, structure and function are similar to humans [26–30]. Through extensive histological characterization of the porcine pancreas across gestation, we report a modest change in β-cell area but a significant expansion of the γ-cells. We observed a reduction in serotonin production within porcine islets contrary to rodent studies. Finally, an adapted protocol for proteomics of formalin fixed paraffin embedded (FFPE) pancreas sections identified solute carrier family 20 member 2 (SLC20A2) and zinc finger CCHC-type containing 7 (ZCCHC7) as significantly downregulated proteins in early pregnancy with potential functions in islet biology. Our results, benchmark the porcine pancreas adaptation to rodent and human studies and highlight the suitability of this model for further studies into gestational pancreas biology.

## Results

### Increase of pancreatic polypeptide cell area in early and late porcine gestation

To map the cellular adaptations of the porcine pancreas during pregnancy, we sampled pig pancreata from non-pregnant (adult) and pregnant pigs at early and late gestation. Our cohort contained a total of 25 animals with similar age distribution (although the late pregnancy animals were slightly older), number of pregnancies and a mean of 167 days post-birth for the non-pregnant control group (Fig. 1A-C). A full description of the cohort is shown on Table 1. As porcine pregnancy spans around 114 days, we classified early and late gestation based on the midgestation timepoint of pregnancy i.e. 57 days of gestation. We focused on the adaptations of the endocrine population of the pancreas by labelling the major hormone populations i.e. insulin, glucagon, somatostatin and pancreatic polypeptide and measured the endocrine area as a fraction of the total area of the pancreas section. Insulin-secreting β-cell area showed a non-significant expansion from early pregnancy suggesting a modest percentage of β-cell increase (Fig. 1D-F,J). No changes were observed in regard to the glucagon-secreting α-cell (Fig. 1D-E,K) or somatostatin-secreting δ-cell area (Fig. 1G-I,L). Interestingly, we observed a robust increase in the pancreatic polypeptide γ-cell area starting in early pregnancy that persisted in late gestational stages (Fig. 1G-I,M). This data suggests a modest β-cell area expansion in porcine pregnancy but a strong increase in the γ-cell area across gestational age.

**Figure 1:**
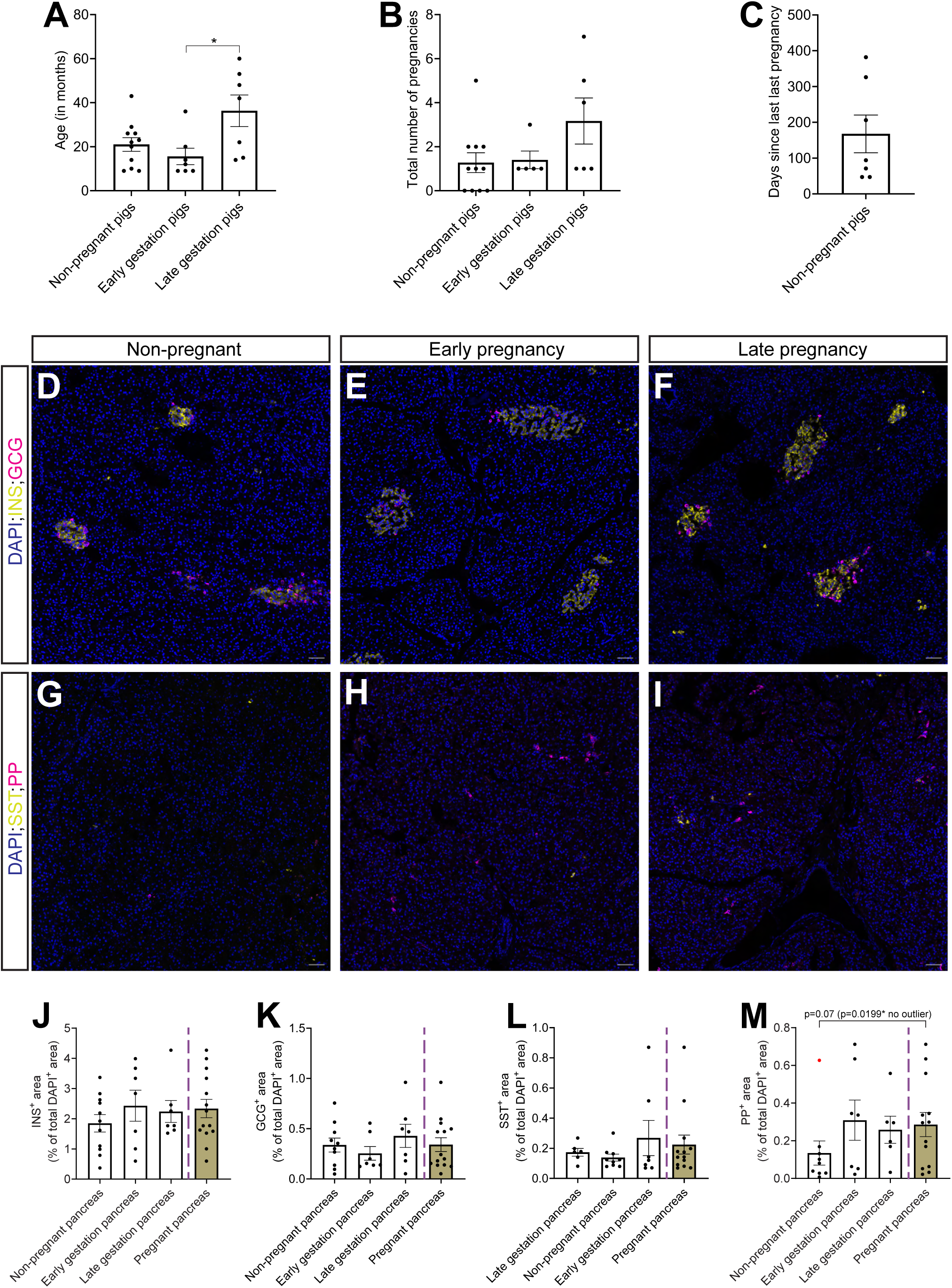
Endocrine cell area expansion in porcine pancreas in early and late gestation. (**A**-**C**) Bar plots showing the age (**A**), total number of pregnancies (**B**) and days since the last pregnancy for the non-pregnant control samples (**C**) of the cohort, when corresponding data was available. Statistical analysis was performed with a Kruskal-Wallis test followed by Dunn’s multiple comparisons test for (**A**). *P=0.0318. (**D**-**F**) Single-plane confocal images of pancreatic sections from non-pregnant (**D**), early pregnancy (**E**) and late pregnancy (**F**) porcine pancreas labelled for INS, GCG and counterstained with DAPI. Scale bar: 50 µm. (**G**-**I**) Single-plane confocal images of pancreatic sections from non-pregnant (**G**), early pregnancy (**H**) and late pregnancy (**I**) porcine pancreas labelled for SST, PP and counterstained with DAPI. Scale bar: 50 µm. (**J**-**M**) Bar plots showing the quantification of the INS (**J**), GCG (**K**), SST (**L**) and PP (**M**) area as a percentage of the total DAPI area per slide and animal. Purple dot line delineates the combined measurement for both early and late pregnancy (Pregnant pancreas, brown bar) from the rest of the graph. The red dot in (**M**) marks the outlier as calculated by ROUT’s test in GraphPad prism. Statistical analysis was performed with a non-parametric Mann-Whitney test. Exact p-values are shown in the graph. *n*= 7-11 (**J**); 7-10 (**K**); 6-9 (**L**);6-9 (**M**). Pregnant pancreas *n* is equal to the sum of the early gestation and late gestation pancreas samples.

**Table 1:** Descriptive characteristics of the pig cohort used in this study. When the information was not available, we marked the entry with an X.

| Pregnancy status at time of pancreas collection | Age (months) | Number of pregnancies | Litter size (only for pregnant animals) | Gestational age (GE) at sacrifice (only for pregnant animals) | Days since last pregnancy (in days - for non-pregnant animals only) |
| --- | --- | --- | --- | --- | --- |
| Non-pregnant 1 | 22 | 2 |  |  | 47 |
| Non-pregnant 2 | 20 | 1 |  |  | 94 |
| Non-pregnant 3 | 26 | 1 |  |  | 326 |
| Non-pregnant 4 | 14 | 0 |  |  |  |
| Non-pregnant 5 | 43 | 5 |  |  | 71 |
| Non-pregnant 6 | 10 | 0 |  |  |  |
| Non-pregnant 7 | 23 | 2 |  |  | 47 |
| Non-pregnant 8 | 9 | 0 |  |  |  |
| Non-pregnant 9 | 9 | 0 |  |  |  |
| Non-pregnant 10 | 29 | 2 |  |  | 206 |
| Non-pregnant 11 | 26 | 1 |  |  | 382 |
| Early pregnancy 1 | 14 | 1 | X | GE22 |  |
| Early pregnancy 2 | 12 | 1 | X | GE35 |  |
| Early pregnancy 3 | 20 | X | 10 | GE54 |  |
| Early pregnancy 4 | 9 | 1 | X | GE32 |  |
| Early pregnancy 5 | 36 | 3 | 10 | GE23 |  |
| Early pregnancy 6 | 9 | 1 | 4 | GE40 |  |
| Early pregnancy 7 | 9 | 1 | X | GE22 |  |
| Late pregnancy 1 | 53 | X | 12 | GE77 |  |
| Late pregnancy 2 | 22 | 1 | X | GE62 |  |
| Late pregnancy 3 | 48 | 5 | 19 | GE110 |  |
| Late pregnancy 4 | 14 | 1 | X | GE64 |  |
| Late pregnancy 5 | 15 | 1 | 16 | GE63 |  |
| Late pregnancy 6 | 60 | 7 | 4 | GE76 |  |
| Late pregnancy 7 | 42 | 4 | X | GE113 |  |

Next, we characterized the cellular mechanisms that could contribute to pancreas and β-cell adaptations. Total pancreas cellular proliferation was induced in early pregnancy marked by Ki67 (Fig. 2A-C,G). Specifically, β-cell proliferation was increased across gestation but the results did not reach statistical significance (Fig. 2D-F,H). No changes in α/β bihormonal cells were observed in the examined sections, suggesting that a prominent transdifferentiation cellular adaptation was unlikely (Fig. 2I-K). Subsequently, we attempted to quantify the number of hormone producing cells within larger ductal structures, as ductal cells have been postulated to be a progenitor source for β-cells. Our analysis showed rare α- or β-cells within the larger ductal structures but with no difference between non-pregnant and pregnant pancreata (only one β-cell within the ductal tree and five α-cells in one ductal structure quantified across all samples). To understand if there is a re-activation of an embryonic differentiation program within ductal cells, we assessed neurogenin 3 (*NEUROG3*) expression by employing the RNAScope methodology. While we observed rare, actively transcribing *NEUROG3* ductal cells in pregnant pancreas, these events were not frequent enough for a meaningful quantification (0-1 cells per section) (Supplementary Fig. 1A-D). Collectively these results correlate with the modest expansion of β-cell area and point to a tissue-wide proliferation induction in early pregnancy.

**Figure 2:**
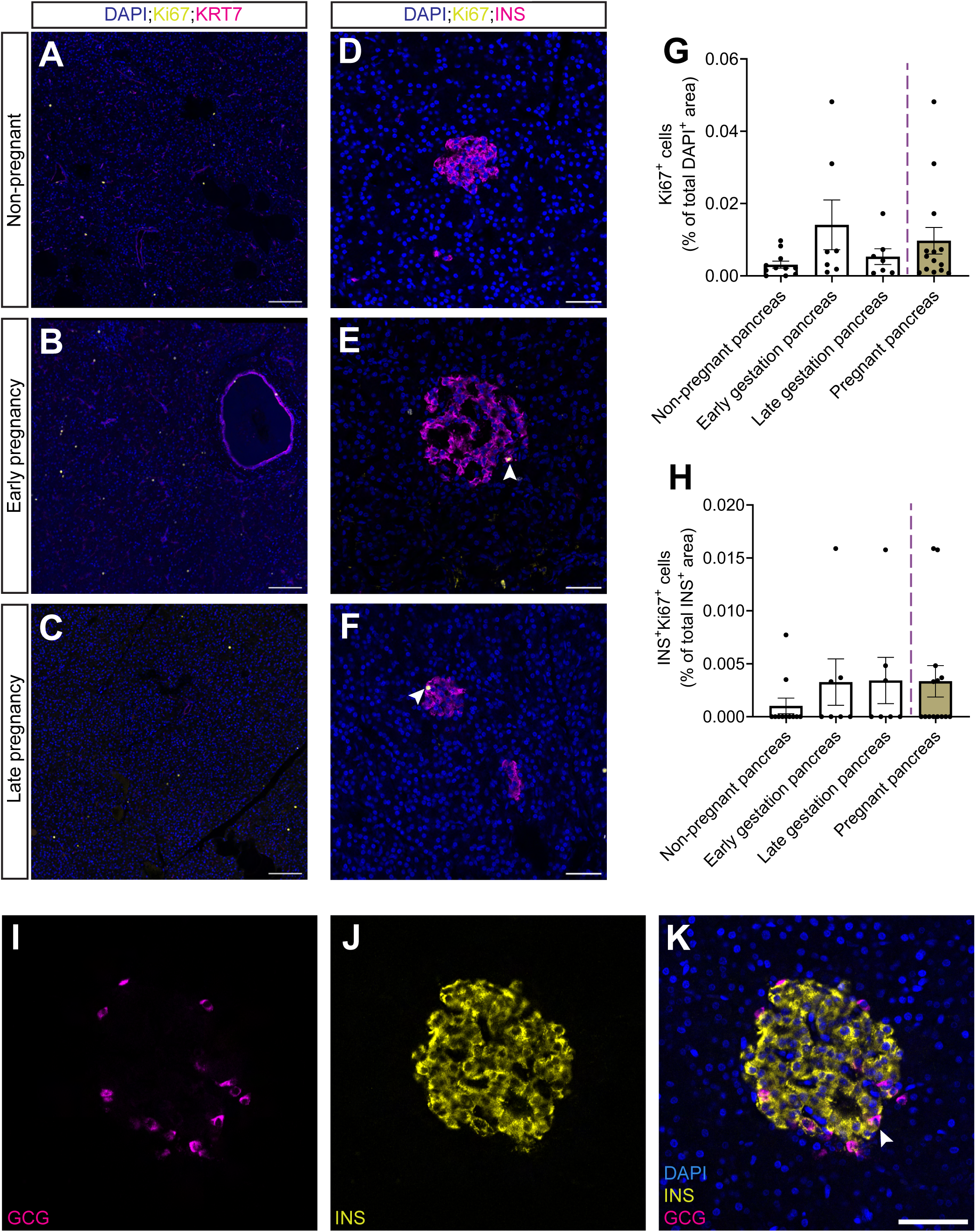
Cellular mechanism of porcine pancreatic adaptations during pregnancy. (**A**-**C**) Single-plane confocal images of pancreatic sections from non-pregnant (**A**), early pregnancy (**B**) and late pregnancy (**C**) porcine pancreas immunostained against Ki67 (proliferation marker), KRT7 (ductal marker) and counterstained with DAPI. Scale bar: 100 µm. (**D**-**F**) Single-plane confocal images of pancreatic sections from non-pregnant (**D**), early pregnancy (**E**) and late pregnancy (**F**) porcine pancreas immunostained against Ki67 (proliferation marker), INS and counterstained with DAPI. Scale bar: 50 µm. (**G**-**H**) Bar plots showing the quantification of the total tissue (**G**) and β-cell (**H**) proliferation percentage across gestation. Purple dot line delineates the combined measurement for both early and late pregnancy (Pregnant pancreas, brown bar) from the rest of the graph. Statistical analysis was performed with a non-parametric Kruskal-Wallis test followed by Dunn’s multiple comparisons test. *n*=7-11 for (**G-H**). (**I**-**K**) Single-plane confocal images of a pancreatic section from a non-pregnant pig pancreas showing the GCG (**I**), INS (**J**) and merged panel (**K**) with DAPI counterstaining. Arrowhead points to a co-expressing GCG/INS bi-hormonal cell. Scale bar: 50 µm. *n*=7-11.

### Serotonin production and uptake is reduced during gestation

Identifying the evolutionary conserved molecular signals that could be implicated in endocrine pancreas adaptation under insulin resistance can lead to new pathways for inducing β-cell regeneration in diabetes. Serotonin signaling has been highlighted as one of the major drivers of pancreatic β-cell adaptations in rodent models of pregnancy. To assess its impact on porcine gestation, we probed the expression levels of serotonin (5-hydroxytryptamine, 5-HT) together with the expression of the rate-limiting enzyme for serotonin production, tryptophan hydroxylase 1 (TPH1). In general, 5-HT was prominent and specific to porcine islets across all tested conditions (Fig. 3A,D,G). Moreover, we discovered few TPH1^+^ cells within islet structures, suggesting some level of 5-HT production within porcine islets (Fig. 3B,E,H). Quantification of the 5-HT^+^ and TPH1^+^ area showed a significant decrease along the gestational progression in porcine pancreas (Fig. 3A-K). Moreover, we assessed the expression levels of the C-JUN transcription factor, which has been postulated to be an effector downstream of prolactin receptor signaling induced in rodent pregnancy [31]. Protein levels and cellular specificity of C-JUN were unchanged between pancreata from non-pregnant and pregnant pigs (Fig. 3L-N). These results indicate that pancreas adaptation on the molecular level can have varying effects between species during pregnancy.

**Figure 3:**
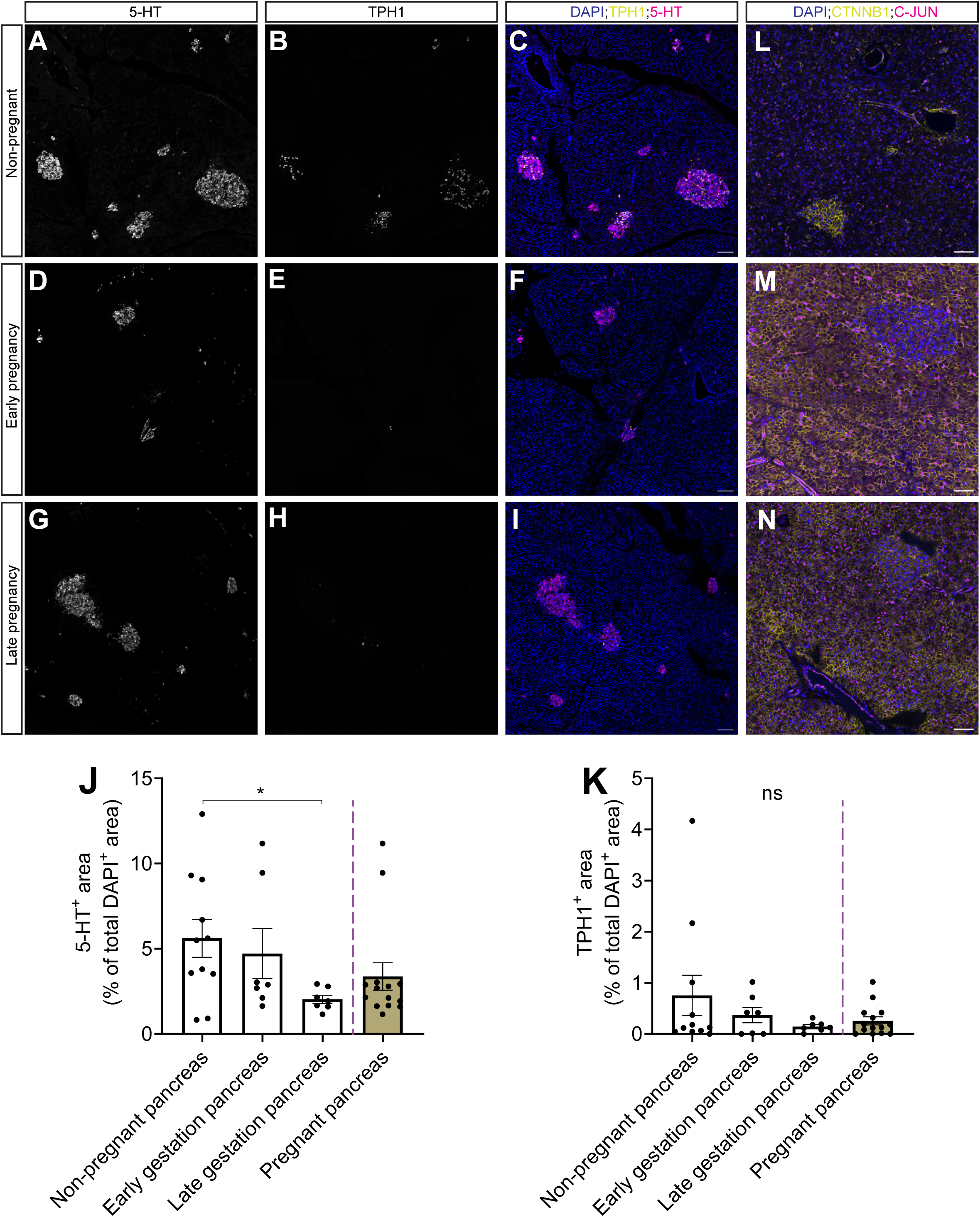
Serotonin production is decreased during pig gestation. (**A**-**I**) Single-plane confocal images of pancreatic sections from non-pregnant (**A-C**), early pregnancy (**D-F**) and late pregnancy (**G-I**) porcine pancreas labelled for 5-HT (**A**,**D**,**G**), TPH1 (**B**,**E**,**H**) and counterstained with DAPI (merge images – **C**,**F**,**I**). Scale bar: 100 µm. (**J**-**K**) Bar plots showing the quantification of the 5-HT (**J**) and TPH1 (**K**) area as percentage of the total section area. Purple dot line delineates the combined measurement for both early and late pregnancy (Pregnant pancreas, brown bar) from the rest of the graph. Statistical analysis was performed with a non-parametric Kruskal-Wallis test followed by Dunn’s multiple comparisons test. *P=0.0460. *n*=7-11 for (**J-K**). (**L**-**N**) Single-plane confocal images of pancreatic sections from non-pregnant (**L**), early pregnancy (**M**) and late pregnancy (**N**) porcine pancreas immunostained against CTNNB1 (epithelial marker), C-JUN and counterstained with DAPI. Scale bar: 50 µm. *n*=2-3 per gestation period.

### Pancreatic proteomics changes in porcine pregnancy

To understand the molecular signals induced in porcine pancreas during pregnancy, we performed proteomics characterization of FFPE pancreas sections across gestation. Briefly, FFPE tissue sections were digested for protein extraction and using mass spectrometry analysis we achieved an average of almost 6000 proteins detected across samples (Fig. 4A). We clustered the samples using Principal Component Analysis (PCA) following imputation but we did not observe a clear separation between the pregnant and non-pregnant tissue, suggesting that the proteome does not undergo extensive molecular remodeling (Fig. 4B). Differential protein analysis identified four proteins to be significantly downregulated in early pregnancy, namely ADAT1, CACFD1, SLC20A2 and ZCCHC7 (Fig. 4C). On the contrary, no significant changes were observed between the non-pregnant and late pregnancy samples (Fig. 4D). Several proteins showed a strong difference between non-pregnant and pregnant samples (Supplementary File 1). For instance, we observed an upregulation in pregnant samples of MCM2 and CENPC, proliferative markers and FAM120b, a protein upregulated in human pregnant pancreata (Fig. 4E-F proteins significant at *pval*<0.05 but not *padj*<0.05 threshold). To validate that our approach could technically detect endocrine-specific proteins given their scarcity in bulk approaches, we measured the endocrine hormone levels in the dataset. All four major hormones were detected in our proteome and interestingly, we observed an increase in pancreatic polypeptide levels across gestation, correlating with the increase in the γ-cell area shown in our histology population (Supplementary Fig. 2A-D). Finally, using our previously published single-cell RNA-Seq dataset of porcine pancreas [32], we explored the expression of differentially altered proteins in the respective cell populations. No cell type enrichment was evident in any of the proteins at mRNA level (Supplementary Fig. 2E-H). These results show a small degree of molecular changes in porcine pancreas during pregnancy, similar to what was recently described in humans [23].

**Figure 4:**
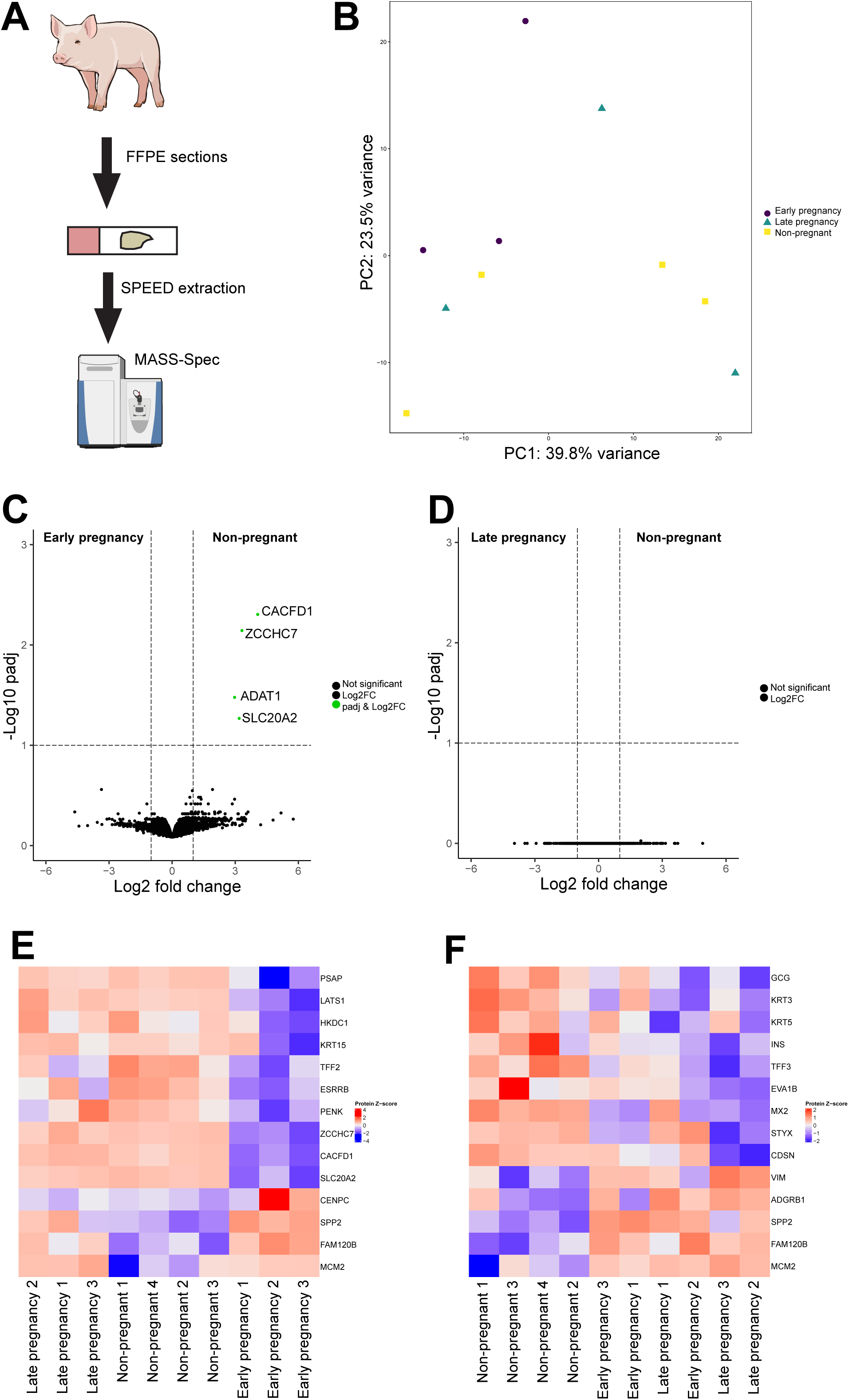
Proteomics characterization of porcine pancreatic adaptations during pregnancy. (**A**) Schema showing the experimental procedure for FFPE-based proteomics characterization of porcine pancreas. (**B**) PCA plot showing the clustering of the different samples following data imputation using the QRILC method. (**C**-**D**) Volcano plots showing the differentially altered proteins comparing the non-pregnant with the early pregnancy (**C**) and the non-pregnant with the late pregnancy (**D**) pancreatic samples. Green dots highlight the significantly regulated proteins at padj <0.1. (**E**-**F**) Heatmaps of the proteomics dataset showing downregulated and upregulated proteins for the non-pregnant versus early pregnancy (**E**) and the non-pregnant versus late pregnancy (**F**). Proteins were shown were changed at pval<0.05.

Subsequently, we assessed the expression level of the differentially expressed proteins to the pancreatic tissue. Two commercially available antibodies gave reproducibly positive signals for SLC20A2 and ZCCHC7 in our porcine sections. Both proteins exhibited an islet-restricted expression pattern with SLC20A2 being specific to the pancreatic δ-cell population, while ZCCHC7 marking all islet cell populations (Fig. 5A,A’,A’’&D,D’,D’’). Correlating with the proteomics results, both proteins exhibited low expression in early pregnancy but partially restored to the non-pregnant expression levels in late gestation (Fig. 5A-F). Since these proteins could mark molecular processes relating to hormone secretion and signaling, we probed their expression during postnatal porcine islet maturation i.e. in pancreas collected directly after birth (postnatal day 3), following weaning (postnatal day 40) and in adolescent pancreas (postnatal day 100). Both proteins were expressed from the early postnatal time window and had similar expression levels in pre- and post-weaning pancreas, a period characterized by extensive metabolic remodeling of endocrine cells (Fig. 5G-L). These results validate SLC20A2 and ZCCHC7 as potential regulators of endocrine cell function during early mammalian pregnancy.

**Figure 5:**
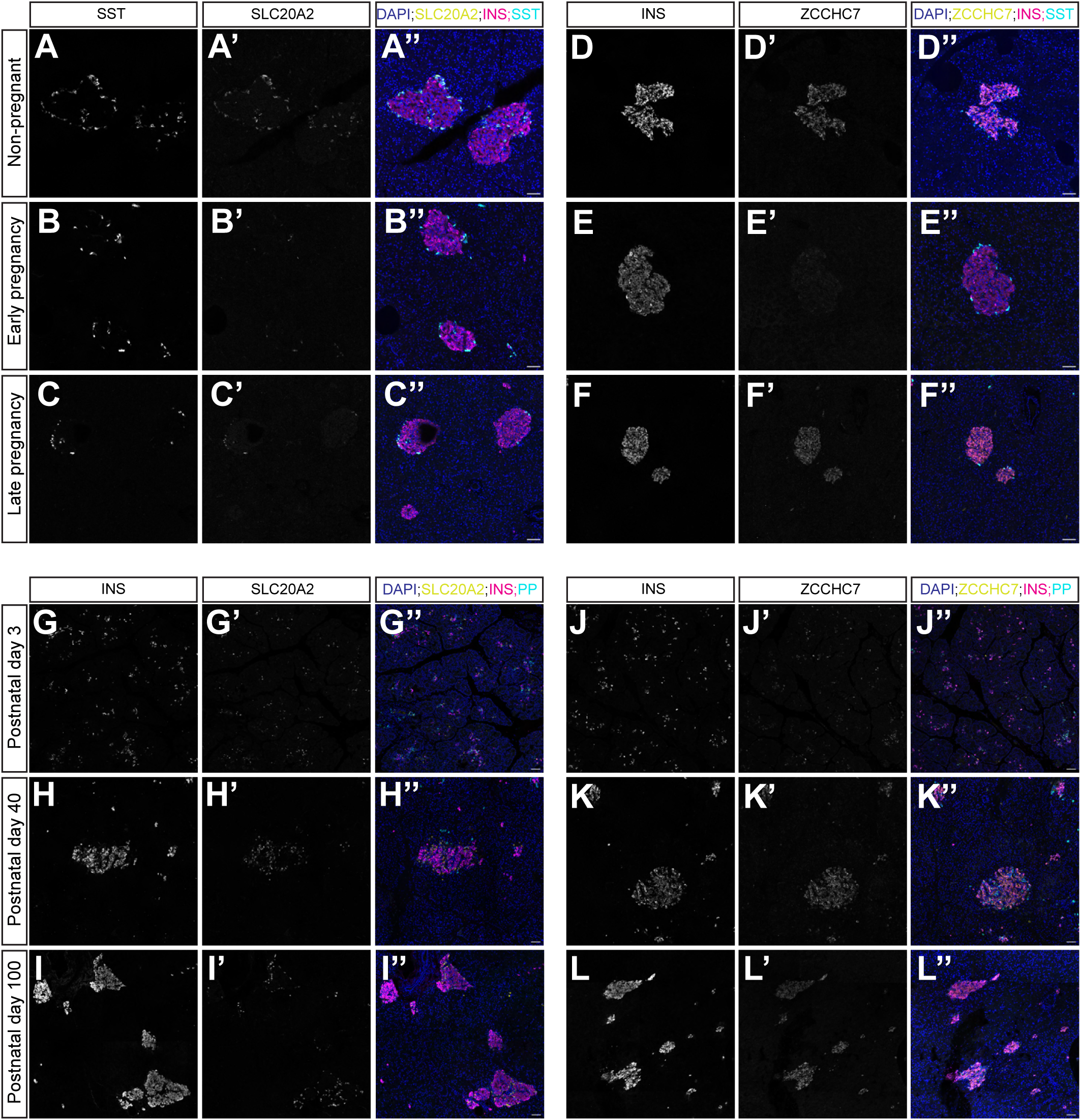
Proteomics validation reveal SLC20A2 and ZCCHC7 as potential regulators of hormone signaling in early pregnancy. (**A**-**F**) Single-plane confocal images of pancreatic sections from non-pregnant (**A-A’’**&**D-D’’**), early pregnancy (**B-B’’**&**E-E’’**) and late pregnancy (**C-C’’**&**F-F’’**) porcine pancreas labelled for SST (**A**-**C**), SLC20A2 (**A’-C’**) and counterstained with DAPI (merge images – **A’’-C’’**), as well as for INS (**D**-**F**), ZCCHC7 (**D’-F’**) and counterstained with DAPI (merge images – **D’’-F’’**). Scale bar: 50 µm. (**G**-**L**) Single-plane confocal images of pancreatic sections from non-pregnant (**G-G’’**&**J-J’’**), early pregnancy (**H-H’’**&**K-K’’**) and late pregnancy (**I-I’’**&**L-L’’**) porcine pancreas labelled for INS (**G-I**), SLC20A2 (**G’-I’**) and counterstained with DAPI (merge images – **G’’-I’’**), as well as for INS (**J**-**L**), ZCCHC7 (**J’-L’**) and counterstained with DAPI (merge images – **J’’-L’’**). Scale bar: 50 µm.

## Discussion

Maternal metabolic adaptations during pregnancy are crucial for ensuring proper progression of fetal development while preventing the establishment of gestational diabetes. Pancreatic adaptations have been mainly studied in rodents, but the translation of these results to mammals with longer gestational period are lacking. To address this gap, we present a comprehensive histological characterization of the pancreatic changes in pig pregnancy. Our model enables us to study the early gestation period, which is not feasible to achieve with human donor samples, in a model with comparable gestational dynamics and metabolism. We report a modest increase of the β-cell area but a significant increase in the γ-cell area along with a significant decrease of the serotonin production along gestation. Molecular insights by FFPE proteomics on pig pancreas tissue across gestation identified SLC20A2 and ZCCHC7 as potential regulators of endocrine biology during gestation. Our results highlight the advantages of the pig model for studying pregnancy related pancreatic adaptations.

The modest β-cell area expansion and proliferation correlates well with the few histological studies performed on human cadaveric pancreata [21–23] and are in direct contrast to the rodent studies. It has been reported that there is an almost four-fold induction of β-cell mass in mice and to our knowledge no study has characterized the γ-cell population, which was increased in pigs [7]. An unresolved question is whether pancreatic adaptation during pregnancy can also occur through pancreatic progenitors, as observed in mice [15]. Transcriptomics characterization of a human ductal xenograft model in pregnant mice proposed some endocrine gene induction but no clear induction of *NEUROG3* [17]. While we observed rare NEUROG3^+^ cells within the ductal epithelium, the absence of suitable lineage-tracing in our porcine study makes our observations rather descriptive. Importantly, we observed a reduction in serotonin synthesis/uptake during pig pregnancy, which contradicts both mouse and human observations[10,11,23]. Few porcine islet cells expressed TPH1, which could argue that the 5-HT abundance could be due to uptake. Human islets have been shown to produce serotonin outside of pregnancy [33], and recent observations have implicated the solute carrier family 18 member a1, which is involved in serotonin signaling, in human endocrine development [34].

Collectively these results highlight species-specific pancreatic adaptations in terms of cellular and molecular signaling. Suitable porcine lineage tracing models can potentially resolve the cellular mechanism of porcine pancreas adaptations with higher resolution in the future and clarify the paracrine regulation based on the γ-cell expansion in early pregnancy.

In terms of molecular signaling, we performed an unbiased profile of the porcine pancreas across gestational ages using FFPE proteomics. The relatively high number of unique protein identifications suggests that our pipeline could robustly identify molecular changes on the protein level. Yet, we could only detect minimal changes on the pancreas proteome, suggesting that the pancreatic adaptations are less than could be expected when compared to rodent transcriptomics data [24,25,35]. Importantly, our FFPE proteomics effect sizes agree with the recent human FFPE proteomics from pancreatic samples of pregnant donors [23] showing the strong correlation between the two species. Despite not reaching significant levels, certain proteins were trending towards similar effect sizes, exemplified by FAM120b, which was upregulated in both porcine and human datasets. In our study, we validated SLC20A2 and ZCCHC7 as two islet-enriched proteins specifically downregulated in early pregnancy. *SLC20A2*, encoding a sodium-phosphate symporter, resides in proximity to a differentially methylated region in type 2 diabetes human islets with open chromatin configuration, pointing to a potential regulation of hormone biology in situations of increased metabolic demand [36]. Not much is known about ZCCHC7, belonging to an RNA metabolism family of proteins [37], in relation to pancreas biology. Nevertheless, increasing the number of biological replicates and coupling them with different modalities will likely provide a more robust molecular understanding of mammalian pregnancy using the pig as a model.

In conclusion, our work demonstrates the similarities between the porcine and human pregnancy adaptations of the pancreas in terms of proteome rewiring and histology. A future comprehensive metabolite and proteome plasma profiling of pig pregnancy will accurately assess its systemic relevance between non-human primate [38] and human global adaptations [39]. Our findings demonstrate that the porcine model can be used to infer meaningful biological insights into mammalian pregnancy and be utilized for getting insights into gestational diabetes mechanisms.

## Materials and Methods

### Pig colony management

Pigs from different transgenic backgrounds (German landrace) were used for this study with a sampling method of convenience, meaning than pancreata were sampled following pig euthanization for other planned experiments. Pigs were maintained at the pathogen-free pig facility of LMU Munich (Center for Innovative Medical Models; www.lmu.de/cimm/). Health assessment was routinely carried out according to FELASA guidelines. Work described in this paper was performed under the Permission No. 55.2-2532.Vet_02-17-136 and 55.2-2532.Vet_02-19-195 approved by the licensing committee from the responsible authority (Government of Upper Bavaria). All experiments were conducted according to the German Animal Welfare Act and Directive 2010/63/EU on the protection of animals used for scientific purposes.

### Porcine pancreas processing and staining

Isolated whole pancreata was cleaned from larger vessels and connective tissue, smaller pieces were cut (2-5 cm) and fixed with 4% paraformaldehyde overnight. After washing three times with PBS 1X, half of the pieces were allocated for cryo-embedding and half were used for paraffin embedding, when sample number was permitting. For cryo-embedding, pancreas was incubated consecutively with 15% and 30% of sucrose solution in PBS 1X overnight and embedded in tissue freezing medium before storing at -80°C, until use. For paraffin embedding, pancreas was dehydrated with a gradient of alcohol before embedding into melted paraffin. Samples were stored at 4°C degrees until further processing.

Both cryo-embedded and paraffin-embedded tissues were sectioned for all stainings described in the process. Cryo-embedded tissue was sectioned at 12 µm thickness with the use of a cryostat (Leica Biosystems) and paraffin-embedded tissue was sectioned at 4 µm thickness with a microtome (Leica Biosystems).

For immunohistochemistry, cryosections were first rehydrated for 30’ with PBS 1X, permeabilized with a solution of 0.1M glycine, 0.3% Triton X-100 in PBS 1X for 10’ followed by blocking solution (3% donkey serum,0.1% bovine serum albumin, 10% fetal calf serum diluted in PBS 1X) and overnight incubation with primary antibodies at 4°C. Following three washes with PBS 1X secondary antibodies were diluted at 1:800 and incubated with the tissue for two hours at room temperature. Following three 10’ washed with PBS 1X, sections were mounted using elvanol and stored at 4°C until imaging. Paraffin-embedded sections were firstly rehydrated with a xylene (2X, 5’ in xylene) followed by ethanol gradient combination (2X in 100% ethanol, 1X in 90% ethanol, 1X in 70% ethanol, 1X in 50% ethanol and 2X in MilliQ water).

Antigen retrieval was used to recover epitopes by microwave heating the sections in sodium citrate pH 6 for 1’ at 780W followed by 14’ at 280W. Sections were let to cool at room temperature for 1h after which the immunostaining protocol was performed as described for cryosections. Primary antibodies used in this study are: a-INS (guinea pig, 1:200, LSBio (BIOZOL), LS-C85862-1), a-INS (rabbit, 1:600, Cell signaling,3014), a-GCG (guinea pig, 1:1000, Takara, M182), a-SST (mouse, 1:400, Santa Cruz, sc-55565), a-SST-PE (1:200, Miltenyi Biotec, 130-127-359), a-PP (goat, 1:200, Sigma Aldrich, SAB2500747), a-KRT7-FITC (1:100, Miltenyi Biotec, 130-115-446), a-CPA1-PE (1:100, MIltenyi Biotec, 130-131-852), a-Ki67 (1:200, rabbit, abcam, ab16667), a-5HT (rabbit, 1:200, Neuromics, RA20080), a-TPH (1:100, sheep, Merck Millipore, AB1541), a-SLC20A2 (1:400, rabbit, Proteintech, 16870734), a-ZCCHC7 (1:200, rabbit, Atlas antibodies, HPA020600), a-ADAT1 (1:100, rabbit, Atlas antibodies, HPA040903 ), a-CACFD1 (1:100, rabbit, Atlas antibodies, ATA-HPA015280-100). Alexa fluorophore conjugated secondary antibodies were used in suitable combinations across the work to label primary antibodies with fluorescence when necessary.

RNAscope in situ hybridization

In situ hybridization was performed using the RNAscope methodology (Advanced Cell Diagnostics) on formalin–fixed, paraffin–embedded (FFPE) tissue sections. Slides were baked at 60 °C for 1 h, deparaffinized in xylene (2 × 15 min), and rehydrated through graded ethanol solutions (100% [2×], 96%, and 70%; 2 min each). Following rehydration, endogenous peroxidase activity was quenched by incubation with hydrogen peroxide solution for 10 min, followed by two washes in distilled water.

Antigen retrieval was performed using a steamer. Slides were pre–incubated in pre–warmed distilled water for 10 s and subsequently incubated with RNAscope Co–Detection Target Retrieval reagent for 30 min. Protease treatment was carried out using RNAscope Manual PretreatPRO reagent for 30 min at 40 °C in the HybEZ™ II oven. RNAscope probe (Ss-NEUROG3-O1, ACD, 498781) was then applied to tissue sections and hybridized for 2 h at 40 °C. Following hybridization, slides were stored in 5× saline sodium citrate (SSC) buffer at 4 °C overnight. On the following day, sequential amplification steps were performed using AMP1, AMP2, and AMP3 reagents according to the manufacturer’s instructions. HRP conjugates corresponding to the respective probes were incubated for 15 min at 40 °C, followed by signal development with the appropriate fluorophores and application of the HRP blocker reagent. For combined RNAscope and immunofluorescence detection, primary and secondary antibodies were applied as described in the relevant section. Slides were mounted using Gold antifade mounting medium and imaged using the Stellaris confocal microscopy system.

### Imaging and image analysis

Images were acquired either with a Nikon SlideScanner to capture the whole tissue area or a Stellaris laser-scanning (Leica Biosystems) confocal microscope for higher resolution images. Whole tissue slices were analyzed with the Fiji software after thresholding the positive areas for each quantification shown in the manuscript.

### Proteomics data acquisition and analysis

For proteomics sample preparation, we de-paraffinized 20 µm thick paraffin-embedded sections from four non-pregnant, three early pregnancy and three late pregnancy samples. Deparaffinization was performed according to the recently described protocol for human pancreas paraffin tissue [23]. Following deparaffinization, the tissue was scraped into a low-protein bind Eppendorf tube and stored at -80°C until further processing.

De-paraffinized material was subjected to the SPEED protocol [40]. After protein quantification using BCA (ThermoFischer Scientific), ten µg per sample were reduced, alkylated and digested in-solution using trypsin and Lys-C, overnight. Peptides were purified as described previously[41] and stored at -20°C until measurement. The MS data were acquired in DIA-PASEF mode on a timsTOF HT mass spectrometer (Bruker, Bremen, DE). Equal amounts of peptides were automatically loaded to the online coupled VanquishNeo (Thermo Fisher Scientific, Dreieich, DE) HPLC system. A Nano-Trap column was used (300-µm inner diameter (ID) × 5 mm), packed with Acclaim PepMap100 C18, 5µm, 100 Å from LC Packings, Sunnyvale, CA, USA, before separation by reversed-phase chromatography (AuroraUltimate 75µm ID × 250 mm, 1.7µm from IonOptics) at 40°C. Peptides were eluted from the column at 250 nL/min using increasing ACN concentration in 0.1% formic acid from 3 to 40% over a 45-min gradient. The DIA-PASEF method covered a mass range from 300 to 1,250 m/z and a mobility range from 0.65 to 1.35 1/ko. Precursor peptides were isolated with 34 equal windows. Collision energy for 0.6 1/ko was set to 20 and for 1.6 1/ko to 59. Estimated cycle time was 1.91 seconds.

DIA files were processed with Spectronaut (Version 20, Biognosys) as direct DIA analysis against a custom-built pig protein database comprised of SwissProt pig proteins combined with non-redundant Trembl sequences, selected for longest sequences per protein entry based on human gene names (16534 sequences; version 1 generated February 2024) using BSG factory settings for Pulsar search. For DIA analysis, default settings were applied. For quantification, precursor filtering was set on Qvalue, with cross run normalization. Quantity MS level was set to MS2; Quantity type was area, and protein quantification is based on the summed-up peptide intensities. PCA plot was calculated on the top 500 most variable proteins. Differential protein analysis was performed with the DEP2 pipeline for comprehensive proteomics data analysis [42]. We imputed the data with the QRILC method and corrected for FDR using Storey’s value, padj<0.1 was considered significant. For the heatmaps, proteins depicted have pval<0.05.

### scRNASeq re-analysis

The normalized values and final annotation from the published dataset (GEO accession: GSE307148 [32]) were used to plot the violin plots using the scanpy suit of tools [43].

### Statistical analysis

Statistical analysis was performed using the GraphPad Prism (version 10.6.1). First samples were tested for normality before employing the statistical tests fit in each analysis and which are mentioned in the figure legends of the work.

## Data availability

The mass spectrometry proteomics data have been deposited to the ProteomeXchange Consortium via the PRIDE [44] partner repository with the dataset identifier PXD077246.

## Acknowledgments

This study was supported by Helmholtz Society and the German Center for Diabetes Research (DZD). CK has received funding from the European Union’s Horizon Europe research and innovation program under the Marie Skłodowska-Curie grant agreement 101108496. MMvDB was supported by the Helmholtz Research School for Diabetes (HRD), which is funded by the Helmholtz Association - Initiative and Networking Fund (IVF).

## Author contributions

C.K. designed the study, performed and analyzed most experiments. S.B. performed histological processing, characterization and imaging of porcine pancreas. C.v.T. performed proteomics data acquisition and analysis. M.M.v.D.B. assisted in sample collection and processing and established image analysis pipelines. D.V. and K.Y. assisted in sample collection, processing and data acquisition E.W. and E.K. performed and supervised pig animal work. H.L. supervised the study and assisted in data analysis. C.K. wrote the first draft of the manuscript and all authors have read and commented on the work.

## Conflict of interest

The authors declare no conflict of interest related to this work.

---

**Supplementary Figure 1:**
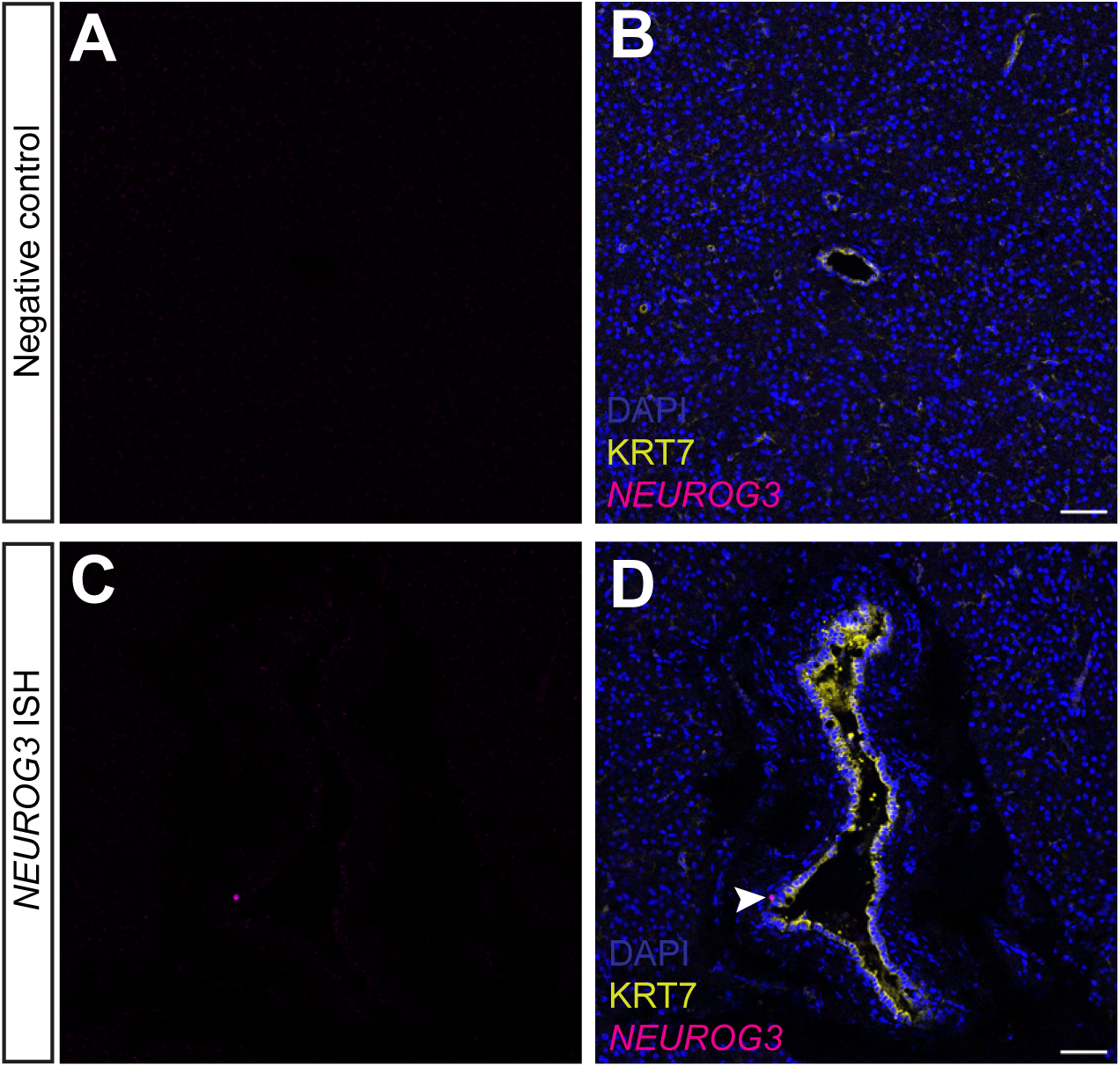
RNA expression levels of *NEUROG3* in porcine pancreas. (**A**-**D**) Single-plane confocal images of the negative control (**A**-**B**) and the NEUROG3 in-situ hybridization (**C**-**D**) results using the RNAScope methodology from an early pregnancy pig pancreas. Slides were immunostained with KRT7 and counterstained with DAPI. Scale bar: 50 µm. *n*=2-3 per gestation period.

**Supplementary Figure 2:**
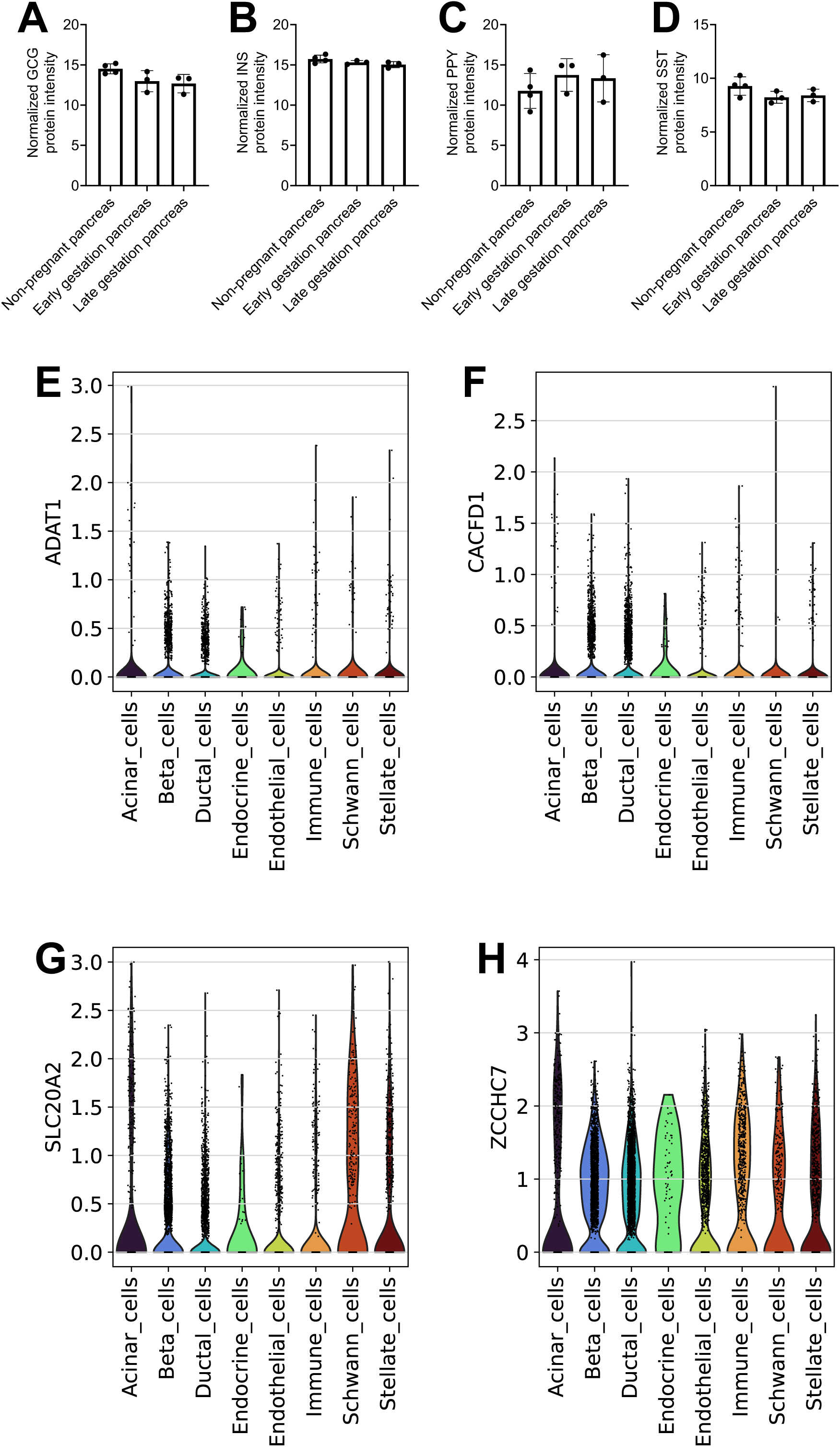
Gene expression levels of differential proteins of the proteomics dataset in pig pancreas. (**A**-**D**) Bar plots show the normalized expression levels of GCG (**A**) and INS (**B**), PPY (**C**) and SST (**D**) in the proteomics dataset. n=3-4. (**E**-**H**) Violin plots showing the mean gene expression levels of *ADAT1* (**E**), *CACFD1* (**F**), *SLC20A2* (**G**) and *ZCCHC7* (**H**) in a scRNA-Seq dataset of porcine pancreas across the annotated cell types.

## Notes

### Competing Interest Statement

The authors have declared no competing interest.

## References

[1] Powe, C.E., Huston Presley, L.P., Locascio, J.J., Catalano, P.M., 2019. Augmented insulin secretory response in early pregnancy. Diabetologia 62(8): 1445–52, Doi: 10.1007/s00125-019-4881-6.

[2] Catalano, P.M., Tyzbir, E.D., Roman, N.M., Amini, S.B., Sims, E.A.H., 1991. Longitudinal changes in insulin release and insulin resistance in nonobese pregnant women. American Journal of Obstetrics and Gynecology 165(6): 1667–72, Doi: 10.1016/0002-9378(91)90012-G.

[3] Jiwani, A., Marseille, E., Lohse, N., Damm, P., Hod, M., Kahn, J.G., 2012. Gestational diabetes mellitus: Results from a survey of country prevalence and practices. Journal of Maternal-Fetal and Neonatal Medicine 25(6): 600–10, Doi: 10.3109/14767058.2011.587921.

[4] Kim, C., Newton, K.M., Knopp, R.H., 2002. Gestational Diabetes and the Incidence of Type 2 Diabetes. Diabetes Care 25(10): 1862–8, Doi: 10.2337/diacare.25.10.1862.

[5] Metzger, B.E., Lowe, L.P., Dyer, A.R., Trimble, E.R., Chaovarindr, U., Coustan, D.R., et al., 2008. Hyperglycemia and adverse pregnancy outcomes. The New England Journal of Medicine 358(19): 1991–2002, Doi: 10.1056/NEJMoa0707943.

[6] Espes, D., Magnusson, L., Caballero-Corbalan, J., Schwarcz, E., Casas, R., Carlsson, P.O., 2022. Pregnancy induces pancreatic insulin secretion in women with long-standing type 1 diabetes. BMJ Open Diabetes Research and Care 10(6), Doi: 10.1136/bmjdrc-2022-002948.

[7] Ruiz-Otero, N., Tessem, J.S., Banerjee, R.R., 2024. Pancreatic islet adaptation in pregnancy and postpartum. Trends in Endocrinology and Metabolism, Doi: 10.1016/j.tem.2024.04.007.

[8] Brelje, T.C., Scharp, D.W., Lacy, P.E., Ogren, L., Talamantes, F., Robertson, M., et al., 1993. Effect of homologous placental lactogens, prolactins, and growth hormones on islet B-cell division and insulin secretion in rat, mouse, and human islets: implication for placental lactogen regulation of islet function during pregnancy. Endocrinology 132(2): 879–87, Doi: 10.1210/endo.132.2.8425500.

[9] Huang, C., Snider, F., Cross, J.C., 2009. Prolactin receptor is required for normal glucose homeostasis and modulation of β-cell mass during pregnancy. Endocrinology 150(4): 1618–26, Doi: 10.1210/en.2008-1003.

[10] Kim, H., Toyofuku, Y., Lynn, F.C., Chak, E., Uchida, T., Mizukami, H., et al., 2010. Serotonin regulates pancreatic beta cell mass during pregnancy. Nature Medicine 16(7): 804–8, Doi: 10.1038/nm.2173.

[11] Moon, J.H., Kim, H., Kim, H., Park, J., Choi, W., Choi, W., et al., 2020. Lactation improves pancreatic β cell mass and function through serotonin production. Science Translational Medicine 12(541), Doi: 10.1126/SCITRANSLMED.AAY0455.

[12] Banerjee, R.R., Cyphert, H.A., Walker, E.M., Chakravarthy, H., Peiris, H., Gu, X., et al., 2016. Gestational diabetes mellitus from inactivation of prolactin receptor and MafB in islet β-cells. Diabetes 65(8): 2331–41, Doi: 10.2337/db15-1527.

[13] Rodriguez, U.A., Socorro, M., Criscimanna, A., Martins, C.P., Mohamed, N., Hu, J., et al., 2021. Conversion of α-Cells to β-Cells in the Postpartum Mouse Pancreas Involves Lgr5 Progeny. Diabetes 70(July): db201059, Doi: 10.2337/db20-1059.

[14] Dubey, V., Tanday, N., Ali, A., Tarasov, A.I., Flatt, P.R., Irwin, N., et al., 2025. Effect of Multiparity on Pregnancy-Induced Islet Adaptation and Cellular Transdifferentiation. Clinical Medicine Insights: Endocrinology and Diabetes 18, Doi: 10.1177/11795514251386124.

[15] Dirice, E., De Jesus, D.F., Kahraman, S., Basile, G., Ng, R.W.S., El Ouaamari, A., et al., 2019. Human duct cells contribute to β cell compensation in insulin resistance. JCI Insight 4(8), Doi: 10.1172/jci.insight.99576.

[16] Toselli, C., Hyslop, C.M., Hughes, M., Natale, D.R., Santamaria, P., Huang, C.T.L., 2014. Contribution of a non-β-cell source to β-cell mass during pregnancy. PLoS ONE 9(6): 1–12, Doi: 10.1371/journal.pone.0100398.

[17] Dirice, E., Basile, G., Kahraman, S., Diegisser, D., Hu, J., Kulkarni, R.N., 2022. Single-nucleus RNA-Seq reveals singular gene signatures of human ductal cells during adaptation to insulin resistance. JCI Insight 7(16), Doi: 10.1172/jci.insight.153877.

[18] Banerjee, R.R., 2018. Piecing together the puzzle of pancreatic islet adaptation in pregnancy. Ann. N.Y. Acad. Sci 1411: 120–39, Doi: 10.1111/nyas.13552.

[19] Fields, A.M., Welle, K., Ho, E.S., Mesaros, C., Susiarjo, M., 2021. Vitamin B6 deficiency disrupts serotonin signaling in pancreatic islets and induces gestational diabetes in mice. Communications Biology 4(1): 333–41, Doi: 10.1038/s42003-021-01900-0.

[20] Assche, F.A., Aerts, L., Prins, F. De., 1978. A MORPHOLOGICAL STUDY OF THE ENDOCRINE PANCREAS IN HUMAN PREGNANCY. BJOG: An International Journal of Obstetrics and Gynaecology 85(11): 818–20, Doi: 10.1111/j.1471-0528.1978.tb15835.x.

[21] Butler, A.E., Cao-Minh, L., Galasso, R., Rizza, R.A., Corradin, A., Cobelli, C., et al., 2010. Adaptive changes in pancreatic beta cell fractional area and beta cell turnover in human pregnancy. Diabetologia 53(10): 2167–76, Doi: 10.1007/s00125-010-1809-6.

[22] Seedat, F., Holden, K., Davis, S., Fischer, R., Bancroft, J., Drydale, E., et al., 2025. A new paradigm of islet adaptations in human pregnancy: insights from immunohistochemistry and proteomics. Nature Communications 16(1): 6687, Doi: 10.1038/s41467-025-61852-5.

[23] Kim, S., Whitener, R.L., Peiris, H., Gu, X., Chang, C.A., Lam, J.Y., et al., 2020. Molecular and genetic regulation of pig pancreatic islet cell development. Development (Cambridge, England), Doi: 10.1101/717090.

[24] Seeberger, K.L., Salama, B.F., Kelly, S., Rosko, M., Castro, C., DesAulniers, J., et al., 2023. Heterogenous expression of endocrine and progenitor cells within the neonatal porcine pancreatic lobes–Implications for neonatal porcine islet xenotransplantation. Xenotransplantation 30(2), Doi: 10.1111/xen.12793.

[25] Renner, S., Blutke, A., Clauss, S., Deeg, C.A., Kemter, E., Merkus, D., et al., 2020. Porcine models for studying complications and organ crosstalk in diabetes mellitus. Cell and Tissue Research 380(2): 341–78, Doi: 10.1007/s00441-019-03158-9.

[26] Yang, K., Spitzer, H., Sterr, M., Hrovatin, K., de la O, S., Zhang, X., et al., 2025. A multimodal cross-species comparison of pancreas development. Nature Communications 16(1): 9355, Doi: 10.1038/s41467-025-64774-4.

[27] Chung, J.-Y., Ruiz-Otero, N., Banerjee, R.R., 2025. c-Jun regulates postpartum β-cell apoptosis and survival downstream of prolactin signaling. Molecular and Cellular Endocrinology 606: 112570, Doi: 10.1016/j.mce.2025.112570.

[28] Karampelias, C., Yang, K., Farkas, F.J., Sterr, M., Molina van Den Bosch, M., Renner, S., et al., 2025. Benchmarking porcine pancreatic ductal organoids for drug screening applications. EMBO Molecular Medicine, Doi: 10.1038/s44321-025-00330-3.

[29] Almaça, J., Molina, J., Menegaz, D., Pronin, A.N., Tamayo, A., Slepak, V., et al., 2016. Human Beta Cells Produce and Release Serotonin to Inhibit Glucagon Secretion from Alpha Cells. Cell Reports 17(12): 3281–91, Doi: 10.1016/j.celrep.2016.11.072.

[30] Zhu, H., Wang, G., Nguyen-Ngoc, K.-V., Kim, D., Miller, M., Goss, G., et al., 2023. Understanding cell fate acquisition in stem-cell-derived pancreatic islets using single-cell multiome-inferred regulomes. Developmental Cell, Doi: 10.1016/j.devcel.2023.03.011.

[31] Chung, J.Y., Ma, Y., Zhang, D., Bickerton, H.H., Stokes, E., Patel, S.B., et al., 2023. Pancreatic islet cell type–specific transcriptomic changes during pregnancy and postpartum. IScience 26(4), Doi: 10.1016/j.isci.2023.106439.

[32] Rieck, S., White, P., Schug, J., Fox, A.J., Smirnova, O., Gao, N., et al., 2009. The transcriptional response of the islet to pregnancy in mice. Molecular Endocrinology 23(10): 1702–12, Doi: 10.1210/me.2009-0144.

[33] Layden, B.T., Durai, V., Newman, M. V., Marinelarena, A.M., Ahn, C.W., Feng, G., et al., 2010. Regulation of pancreatic islet gene expression in mouse islets by pregnancy. Journal of Endocrinology 207(3): 265–79, Doi: 10.1677/JOE-10-0298.

[34] Bysani, M., Agren, R., Davegårdh, C., Volkov, P., Rönn, T., Unneberg, P., et al., 2019. ATAC-seq reveals alterations in open chromatin in pancreatic islets from subjects with type 2 diabetes. Scientific Reports 9(1), Doi: 10.1038/s41598-019-44076-8.

[35] Wang, Y., Yu, Y., Pang, Y., Yu, H., Zhang, W., Zhao, X., et al., 2021. The distinct roles of zinc finger CCHC-type (ZCCHC) superfamily proteins in the regulation of RNA metabolism. RNA Biology 18(12): 2107–26, Doi: 10.1080/15476286.2021.1909320.

[36] Yu, D., Wan, H., Tong, C., Guang, L., Chen, G., Su, J., et al., 2024. A multi-tissue metabolome atlas of primate pregnancy. Cell 187(3): 764–781.e14, Doi: 10.1016/j.cell.2023.11.043.

[37] Liang, L., Rasmussen, M.L.H., Piening, B., Shen, X., Chen, S., Röst, H., et al., 2020. Metabolic Dynamics and Prediction of Gestational Age and Time to Delivery in Pregnant Women. Cell 181(7): 1680–1692.e15, Doi: 10.1016/j.cell.2020.05.002.

[38] Doellinger, J., Schneider, A., Hoeller, M., Lasch, P., 2020. Sample preparation by easy extraction and digestion (SPEED) - A universal, rapid, and detergent-free protocol for proteomics based on acid extraction. Molecular and Cellular Proteomics 19(1): 209–22, Doi: 10.1074/mcp.TIR119.001616.

[39] Kulak, N.A., Pichler, G., Paron, I., Nagaraj, N., Mann, M., 2014. Minimal, encapsulated proteomic-sample processing applied to copy-number estimation in eukaryotic cells. Nature Methods 11(3): 319–24, Doi: 10.1038/nmeth.2834.

[40] Feng, Z., Fang, P., Zheng, H., Zhang, X., 2023. DEP2: an upgraded comprehensive analysis toolkit for quantitative proteomics data. Bioinformatics 39(8), Doi: 10.1093/bioinformatics/btad526.

[41] Wolf, A., Angerer, P., Theis, F., 2018. SCANPY: large-scale single-cell gene expression data analysis. Genome Biology 19(15): 2926–34, Doi: 10.1111/1462-2920.13787.

[42] Perez-Riverol, Y., Bandla, C., Kundu, D.J., Kamatchinathan, S., Bai, J., Hewapathirana, S., et al., 2025. The PRIDE database at 20 years: 2025 update. Nucleic Acids Research 53(D1): D543–53, Doi: 10.1093/nar/gkae1011.

